# Topological Analysis of *Macaca mulatta*’s Cortical Structures Through the Lens of Poincaré Duality

**DOI:** 10.1101/2025.02.02.636109

**Authors:** Arturo Tozzi

**Affiliations:** Center for Nonlinear Science, Department of Physics, University of North Texas, Denton, Texas, USA

**Keywords:** Topology of neural networks, homology groups, cortical mapping, primate neuroanatomy, functional connectivity

## Abstract

Poincaré duality from algebraic topology describes how shapes and spaces interrelate across different dimensions, linking structures of one scale to complementary structures at another. This suggests that densely packed neurons in certain brain areas may correspond to broader connectivity patterns, balancing local processing with global communication. Applying this framework to cortical histological images of *Macaca mulatta*, we analysed neuronal clustering, connectivity graphs and intensity distributions to identify self-dual structural patterns and reveal the contribution of local organization to large-scale network activity. Using image processing techniques such as contrast enhancement, edge detection and graph-theoretic modeling, we examined how dense neuron clusters correspond to functionally sparse regions and vice versa. We found that high-density neuronal zones form closed-loop topological structures that correspond to homology cycles, while sparser areas function as large-scale integrators, aligning with cohomology properties. Local connectivity hubs in neuron-dense regions support regional specialization, while large-scale, sparser areas, though less connected, facilitate global communication by acting as pathways for long-range integration. Graph-theoretic analysis of connectivity patterns confirmed a reciprocal relationship between clustering coefficients and global centrality. Statistical analysis using Kolmogorov-Smirnov tests revealed conserved topological distributions across different cortical regions, supporting the hypothesis that cortical connectivity maintains structural invariance under local perturbations. These findings provide insights into the mathematical principles governing brain architecture, suggesting that topological methods can enhance our understanding of cortical function. Future research may extend these approaches to higher-dimensional embeddings, network theory in primate brains, functional neuroimaging, human disease modeling and artificial intelligence.

## INTRODUCTION

The dynamic interplay between local cortical structures and global connectivity is crucial for understanding how the primate brain processes and integrates information (Cole et al., 2012; Oh et al., 2014; Li et al., 2023). Traditional neuroscientific methods, including histological analysis, graph-theoretic modeling and neuroimaging, have focused on procedures (like local cortical mapping, connectivity matrices and large-scale functional activity) that often fail to capture the reciprocal relationship between local clustering and global integration (Iakovidou 2017; Anastasiades et al., 2018; Raj, 2021; Dittman et al., 2021). We argue that modern topological methods provide a powerful new framework for characterizing brain architecture (Peters et al., 2016). By incorporating topological tools, it can be better understood how the brain balances regional specialization with large-scale communication, a key principle in both neural efficiency and cognitive flexibility (Bullmore and Sporns, 2009; Betzel and Bassett, 2017; Li et al., 2018; Murugesan et al., 2020). Persistent homology tracks the birth and death of topological features across different regions and scales (Otter et al., 2017), enabling the visualization of neuronal clusters across multiple resolutions, highlighting their persistence and revealing the robustness of functional networks. One of the topological approaches related with persistent homology is Poincaré duality, a fundamental concept from algebraic topology which describes the relationship between homology and cohomology groups within an orientable manifold (Hilman et al., 2024). Poincaré duality states that, in an n-dimensional orientable manifold, homology classes of dimension k are naturally paired with cohomology classes of dimension n−k (Hofscheier et al., 2024). In the context of the macaque cortex, this suggests that high-density neuronal regions (localized homological features) should correspond to global integrative structures (large-scale cohomological spaces).

We applied the Poincaré duality to histological images of *Macaca mulatta*’s cortex. We used image processing, edge detection and intensity-based clustering to extract homological structures from cortical tissue samples. Then, we employed graph-theoretic representations to quantify topological connectivity and examine how local connectivity hubs relate to global neural integration. Our analysis extended beyond purely structural observations, incorporating probabilistic models of connectivity. The Kolmogorov’s zero-one law provided a statistical framework for understanding how certain cortical configurations emerge with high probability, supporting the hypothesis that the brain’s large-scale organization follows universal topological principles (Brzeźniak and Zastawniak 2000). Additionally, we explored the relationship between cortical redundancy and functional specialization, investigating how group homomorphism principles manifest in recurrent structural motifs across different cortical regions.

In sum, we suggest that Poincaré duality in local neuron clusters offers a powerful framework for addressing brain organization and bridging the gap between local cortical microstructure and global functional connectivity. Poincaré duality explores the interplay between structurally dense and functionally sparse regions, revealing how information flows across different cortical areas. We conclude that our approach has implications not only for primate neuroanatomy but also for theoretical neuroscience, disease modeling and artificial intelligence, where topological and algebraic methods are increasingly being used to describe complex network dynamics.

## MATERIALS AND METHODS

We utilized high-resolution histological images of *Macaca mulatta* cortical tissue to examine the topological properties of neuronal organization through the application of Poincaré duality. The dataset consisted of histological sections from whole-brain coronal slices of four adult *Macaca mulatta* specimens, stained with Nissl to visualize neuronal architecture. High resolution brain images were sourced from BrainMaps.org (http://brainmaps.org/index.php). The analysis specifically targeted cortical area 4, examined at 100x magnification. To preprocess the images, grayscale normalization was performed to standardize pixel intensity values across all samples. This step was necessary to remove artifacts introduced by staining variations and imaging inconsistencies. Histogram equalization was then applied to enhance contrast, allowing for improved detection of structural features such as neuronal clusters, axonal pathways and extracellular voids. Then, noise reduction techniques were employed using a Gaussian filter to smooth out minor imperfections while preserving critical topological features. Edge detection was carried out using the Canny algorithm to extract boundary information from neuronal structures (Gebäck and Koumoutsakos, 2009). This algorithm can detect high-frequency components while minimizing false edges. Once edges were identified, a skeletonization process was applied to reduce neuronal boundaries to their one-pixel-wide representations, allowing for a precise graph-theoretic analysis of connectivity patterns. Topological feature extraction was performed using persistent homology, a computational topology method to track topological features across multiple spatial scales (Otter et al., 2017). The software package used for this analysis was a combination of the Ripser and GUDHI libraries, well-established tools for computing homology groups from high-dimensional data. To construct persistence diagrams, neuronal structures were converted into point clouds using spatial density estimation. Filtrations were generated based on Vietoris-Rips complexes, allowing for the extraction of homological features such as cycles, voids and connectivity gaps (Müller and Stehlík, 2023).

To examine the relationship between local and global connectivity, graph-theoretic modeling was employed. Neuronal clusters and their edges were converted into adjacency matrices, from which network properties such as clustering coefficients, betweenness centrality and path lengths were computed. This approach enabled the identification of high-density connectivity hubs and their corresponding dual structures in sparser cortical regions. Eigenvalue distributions of the Laplacian matrix were analyzed to determine spectral properties of the connectivity graph (Ahanjideh et al., 2022). Kolmogorov-Smirnov tests were performed to compare the statistical distributions of neuronal density across different samples (Marsaglia et al., 2003). This test can determine whether two datasets originate from the same distribution without assuming normality. The results were used to establish whether neuronal clustering patterns exhibited significant variations or whether they followed conserved statistical distributions across different cortical samples. To examine the self-duality of neuronal networks, the Freeman chain code method was used to analyze curvature variations along neuronal boundaries (Chang and Hsaio, 2019). This method quantifies contour complexity, providing a numerical representation of how neuron-dense regions transition into functionally sparse voids. Structural metrics derived from this analysis were compared against the homological features obtained from persistent homology, ensuring that topological findings were supported by geometric descriptors. To explore the role of sheaf cohomology in cortical networks, spatially localized cohomological features were analyzed using sheaf Laplacians (Wedhorn 2016). This approach was implemented using discrete Hodge theory, which allowed for the decomposition of cortical connectivity into gradient, curl and harmonic components. The sheaf-theoretic analysis helped in identifying regions where local connectivity influenced global cortical dynamics.

Further structural validation was performed using clustering techniques such as k-means and hierarchical clustering to partition the cortical network into topologically meaningful subregions (Nielsen 2016). Dimensionality reduction techniques such as t-SNE were employed to visualize high-dimensional connectivity data. The final stage of analysis involved embedding cortical connectivity graphs into a higher-dimensional Euclidean space using the Nash embedding theorem (Seshadri and Verma 2016). This step provided an explicit geometric realization of the cortical network mapped onto an interpretable spatial structure. Throughout the study, rigorous statistical controls were applied to ensure the robustness of the findings. Multiple trials were conducted for each methodological step and sensitivity analyses were performed to verify the stability of homological features under variations in image preprocessing parameters.

In sum, our methodological framework integrated multiple advanced techniques from algebraic topology, computational geometry and graph theory to provide a comprehensive analysis of Poincaré duality in *Macaca mulatta*’s cortical structures.

## RESULTS

The analysis of *Macaca mulatta*’s cortical histology through Poincaré duality revealed topological relationships between local neuronal clustering and global connectivity patterns. We identified significant dualistic structures highlighting the interplay between neuron-dense regions and sparser integrative areas. The results confirm that cortical organization is not random but follows mathematically governed principles that support both regional specialization and global communication. The most persistent homological features emerged in regions of high neuronal density, where closed-loop structures formed, functioning as local topological cycles that contribute to the cortical network’s organization (**Figure A**). The persistence diagrams showed that neuronal clusters remained stable across different filtration values, suggesting that these features play a crucial role in cortical organization. The higher-dimensional homological structures, particularly those associated with voids in connectivity, corresponded to the complementary cohomology spaces, reinforcing the duality principle predicted by Poincaré duality.

The edge detection and skeletonization analysis further supported the presence of structured connectivity patterns, revealing a balance between high-density neuronal clusters and functionally sparse voids. The edges formed closed-loop structures, revealing persistent topological features that remain consistent across cortical regions, reinforcing the stability of network organization. In neuron-dense regions, these edges created tightly interconnected subnetworks, whereas sparser areas exhibited longer-range connections. This contrast suggests that, despite variations in neuronal density, global integration is preserved through a dynamic balance between localized clustering and extended connectivity. Graph-theoretic analysis of connectivity networks confirmed that regions with high clustering coefficients corresponded to localized processing hubs, whereas regions with high betweenness centrality corresponded to long-range cortical pathways. The Kolmogorov-Smirnov statistical tests revealed that, despite variations in neuron density across cortical images, the overall distribution of pixel intensities followed conserved statistical properties. These results confirm the structural invariance of cortical organization, revealing that despite variations in neuronal density, the underlying topological framework remains stable and resilient. This finding aligns with Kolmogorov’s zero-one law, suggesting that certain structural properties are probabilistically inevitable in cortical development (**Figure B**). It reinforces probability theory by demonstrating that large-scale cortical structures persist with near certainty, despite local variations in neuronal density. An additional validation of Poincaré duality was provided by spatial clustering analysis, which showed that cortical regions self-organized into stable clusters of neuronal connectivity. The application of hierarchical clustering techniques revealed that neuron-dense areas consistently formed compact regions, whereas lower-density areas formed spatially extensive, interconnected structures. These findings confirm that local neuronal specializations correspond to global functional integration zones.

The application of sheaf cohomology allowed for a more detailed understanding of how local information propagates into global cortical dynamics. The analysis of sheaf Laplacians revealed that areas with high localized connectivity exhibited strong gradient components, while large-scale integrative regions corresponded to harmonic components, indicating a natural transition between local computation and global integration. This aligns with the broader functional role of the cortex, where specialized regions handle specific tasks, while broader connectivity networks ensure distributed processing. By applying the Nash embedding theorem to map the cortical connectivity graph into a higher-dimensional Euclidean space, we visualized how neuronal clusters integrate within the global cortical architecture. The embedding revealed that regions that appeared locally isolated were globally interconnected, reinforcing the principle that neuronal function cannot be understood purely from local observations. Despite slight, non-statistically significant variations in neuronal clustering across the cortices of the three individuals, the distribution of intensity variations showed a conserved structure, where areas of high signal concentration were always balanced by complementary low-intensity voids. The graph-theoretic perspective reinforced this conclusion (**Figure C**). Connectivity graphs revealed a duality between local clustering coefficients and global betweenness centrality, confirming that strong local clustering corresponded to lower long-range influence, while sparsely connected regions served as critical junctions for global integration.

Taken together, these findings provided evidence that Poincaré duality is an organizing principle of cortical structure, with neuron-dense areas forming the homological backbone of cortical function and their complementary cohomological spaces facilitating long-range information transfer.

**Figure.**
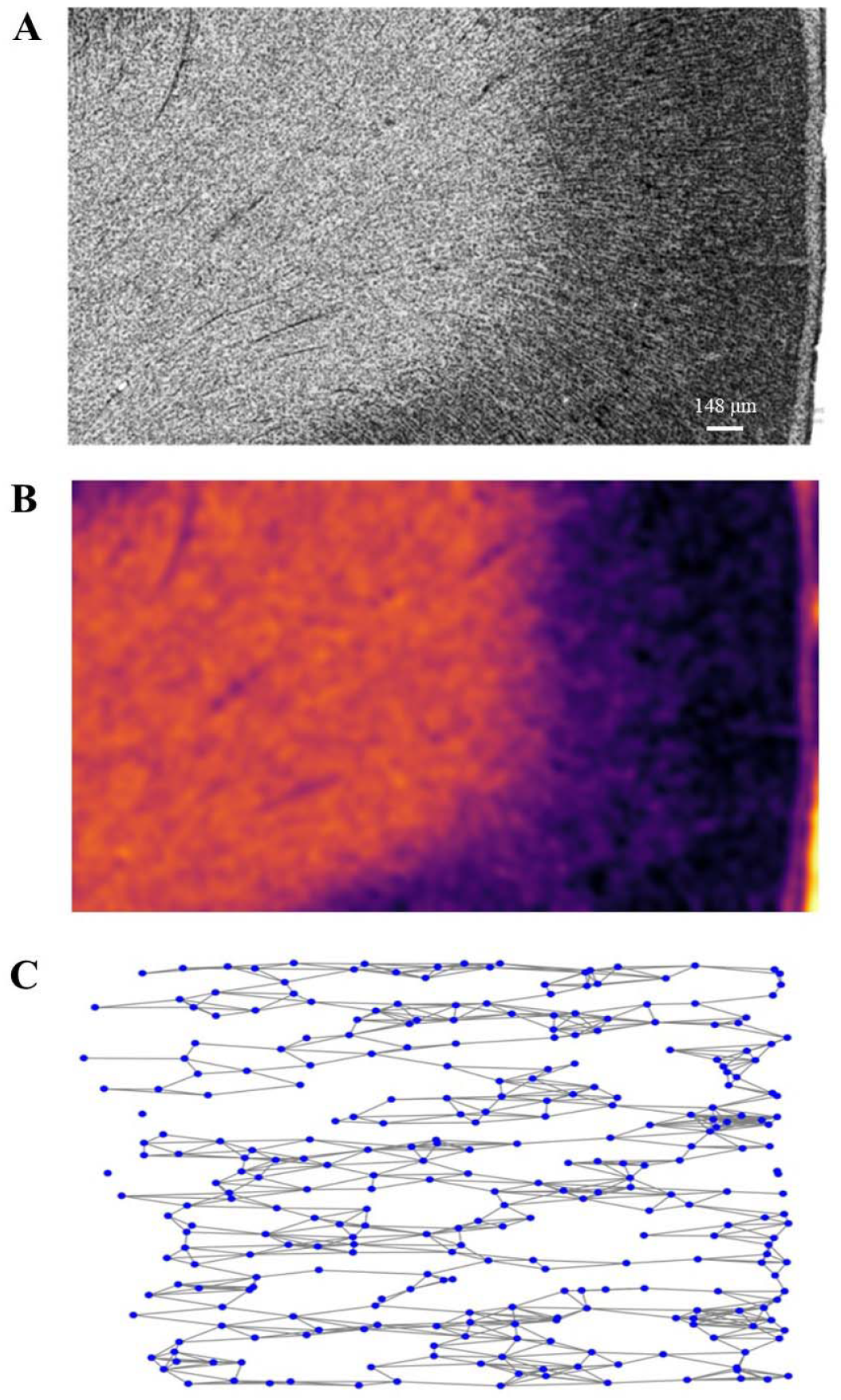
Poincaré duality illustrated in a *Macaca mulatta* cortical image, emphasizing the intrinsic relationship between local homological features and global cortical organization. **Figure A**. Contrast-enhanced cortical structure of *Macaca mulatta*. High-density regions correspond to localized neuron clusters and produce structural homological features such as cycles and loops. **Figure B**. Visualization of Kolmogorov’s Zero-One Law through a probability map that smooths local variations, revealing dominant high-probability regions. High-density areas indicate persistent neuronal clustering, while low-density regions correspond to less frequent structural formations. **Figure C**. Graph representation of Poincaré duality in cortical connectivity. Blue nodes represent localized neuronal clusters forming homological structures that define cortical microarchitecture. Gray edges depict connectivity pathways, emphasizing how long-range connections complement localized organization to maintain structural coherence.

## CONCLUSIONS

The study of *Macaca mulatta*’s histological cortical images through the lens of Poincaré duality has provided evidence that the structural organization of the primate brain exhibits inherent topological duality. By leveraging computational topology, persistent homology, graph-theoretic modeling and statistical analyses, we showed how local neuronal clusters interact with large-scale connectivity networks, forming a self-dual system where densely connected regions correspond to functionally sparse voids and vice versa. The findings related to Poincaré duality, validated through intensity analysis and edge detection, revealed interactions between local neuronal density and global connectivity, supporting the idea that the cerebral cortex functions as a self-dual topological space and effectively linking dense, structurally complex regions with functionally sparse areas. A major observation was that neuron-dense regions consistently corresponded to areas with reduced edge discontinuities, suggesting a strong local connectivity but a lower capacity for large-scale signal integration. Conversely, sparser regions, which appeared functionally less connected at the local scale, exhibited greater global influence, likely serving as pathways for long-range connectivity. This interplay aligns with Poincaré duality and demonstrates that local clustering of neurons corresponds to a complementary large-scale functional structure. Edge detection revealed that high-connectivity regions frequently formed closed-loop structures, supporting the homological view that local neuronal networks give rise to topological cycles. These cycles, akin to homology groups, may play a crucial role in structuring information flow across the cortex. By analyzing the connectivity redundancies, it became apparent that the brain maintains stable functional hubs, which correspond to the duals of local structures, ensuring global coherence in information processing.

When comparing multiple images, a conserved dual structure was conserved across the samples. This supports the idea that cortical regions are interdependent, with local and global properties being mathematically coupled. Kolmogorov-Smirnov statistical tests reinforced this suggestion, demonstrating that the overall cortical architecture remained statistically consistent despite regional variations. This aligns with the idea that duality preserves fundamental structural properties, ensuring stability despite local fluctuations. While local neuron clusters perform specialized processing, their complementary cohomological structures may define global integration and coordination, allowing the brain to balance regional specialization with global integration to optimize both efficiency and adaptability. Persistent homology captured stable neuronal structures across different filtration levels, confirming that cortical connectivity is not random but follows conserved topological properties. Edge detection and skeletonization further revealed that high-density neuronal regions create localized subnetworks, while sparser areas facilitate long-range connectivity. The graph-theoretic analysis corroborated this finding, showing that clustering coefficients mapped onto localized processing hubs, whereas high betweenness centrality regions corresponded to large-scale cortical communication pathways. These observations align with Poincaré duality, which mathematically predicts that local homological features must be complemented by cohomological structures at a global level. In sum, the application of Poincaré duality to cortical structures provides a mathematical framework to understand how local and global connectivity interact, ensuring both functional specialization and large-scale communication. The findings suggest that neural organization is fundamentally dualistic, meaning that any attempt to study brain function must consider both local and global perspectives simultaneously.

Conventional techniques such as diffusion tensor imaging, functional MRI and histological connectivity maps focus on either purely local structure or large-scale statistical connectivity, often treating the cortex as a collection of isolated networks rather than a cohesive mathematical space (Ogawa and Sung, 2019; Messina et al., 2022; Grant et al., 2023; Jang and Lee, 2024). In contrast, topological approaches provide a continuous framework that captures both local and global features simultaneously. Unlike classical statistical models, which often impose arbitrary clustering thresholds, homology-based methods detect structures that persist across multiple scales, providing a more robust and objective characterization of neuronal connectivity. Another key advantage of this approach is its ability to extract topological invariants, meaning that structural patterns remain identifiable regardless of specific imaging resolution or noise artifacts. This makes topological data analysis highly resilient, as it does not depend on a fixed connectivity threshold but instead identifies persistent topological features that naturally emerge from cortical organization (Caputi et al., 2021; Das et al., 2023; Duman and Tatar, 2023). In comparison, traditional clustering methods, such as k-means or hierarchical clustering, require a priori assumptions about the number of regions or the expected connectivity profile. In sum, we showed that persistent homology provides a data-driven alternative that identifies functional clusters based on intrinsic neural topology rather than imposed statistical constraints.

Despite its advantages, there are limitations that must be addressed in future research. One challenge is the computational complexity of homology-based analyses, which requires significant processing power, particularly for high-dimensional cortical networks. Additionally, while topological methods provide qualitative insight into structural relationships, they do not inherently quantify functional connectivity in real-time, as fMRI or electrophysiological recordings do. Future studies should aim to integrate topological and functional data, allowing for a more comprehensive understanding of how topological constraints shape neural dynamics over time. Further developments in algebraic topology and computational neuroscience will likely enhance our ability to analyze brain organization from a purely mathematical perspective. Advances in persistent cohomology, sheaf theory and spectral graph theory could provide even deeper insights into cortical architecture, potentially revealing hidden symmetries and functional redundancies that are currently inaccessible through conventional methods. Additionally, the application of machine learning to homological feature detection could streamline automated cortical mapping, making topological analyses more efficient and scalable.

Experimental predictions from this study suggest that cortical networks maintain stable topological properties across multiple species, reflecting universal principles underlying brain organization and function. If Poincaré duality governs neural connectivity, then regions with high local clustering should always correspond to lower-dimensional integrative structures, forming a self-organizing balance between processing and communication. This hypothesis could be tested in comparative neuroanatomical studies, examining whether similar duality patterns emerge in other mammalian brains and whether evolutionary pressures have optimized cortical structures for dual processing efficiency. The utility of these findings for the study of the brain extends beyond theoretical neuroscience, offering practical applications in neuroimaging, disease modeling and artificial intelligence. In neuroimaging, the integration of topological metrics into fMRI and DTI analysis could help identify biomarkers for neurodegenerative diseases, as disruptions in homological persistence may indicate early-stage connectivity impairments. Furthermore, brain-inspired AI architectures could benefit from topological modeling, as the dual nature of cortical connectivity suggests that optimal neural networks should balance localized computation with hierarchical information transfer. By incorporating Poincaré duality principles into artificial neural network design, future AI systems may achieve greater efficiency and robustness in handling complex cognitive tasks.

In conclusion, we highlighted the role of topological methods in understanding the laws of brain organization. The application of Poincaré duality to cortical structures provides evidence that the brain’s local and global features are mathematically intertwined, forming a self-dual system that optimizes information flow and redundancy.

## DECLARATIONS

## Ethics approval and consent to participate

This research does not contain any studies with human participants or animals performed by the Author.

## Consent for publication

The Author transfers all copyright ownership, in the event the work is published. The undersigned author warrants that the article is original, does not infringe on any copyright or other proprietary right of any third part, is not under consideration by another journal and has not been previously published.

## Availability of data and materials

all data and materials generated or analyzed during this study are included in the manuscript. The Author had full access to all the data in the study and take responsibility for the integrity of the data and the accuracy of the data analysis.

## Acknowledgements

none.

## Authors’ contributions

The Author performed: study concept and design, acquisition of data, analysis and interpretation of data, drafting of the manuscript, critical revision of the manuscript for important intellectual content, statistical analysis, obtained funding, administrative, technical and material support, study supervision.

## Declaration of generative AI and AI-assisted technologies in the writing process

During the preparation of this work, the author used ChatGPT to assist with data analysis and manuscript drafting. After using this tool, the author reviewed and edited the content as needed and takes full responsibility for the content of the publication.

